# Microglia-mediated synaptic pruning in the nucleus accumbens during adolescence: A preliminary study of the proteomic consequences and putative female-specific pruning target

**DOI:** 10.1101/2023.05.02.539121

**Authors:** J. M. Kirkland, Ishan Patel, Ashley M. Kopec

## Abstract

Adolescence is a period of copious neural development, particularly in the ‘reward’ circuitry of the brain, and reward-related behavioral development, including social development. One neurodevelopmental mechanism that appears to be common across brain regions and developmental periods is the requirement for synaptic pruning to produce mature neural communication and circuits. We published that microglia-C3-mediated synaptic pruning also occurs in the nucleus accumbens (NAc) reward region during adolescence to mediate social development in male and female rats. However, both the adolescent stage in which microglial pruning occurred, and the synaptic pruning target, were sex specific. NAc pruning occurred between early and mid-adolescence in male rats to eliminate dopamine D1 receptors (D1rs), and between pre- and early adolescence in female rats (P20-30) to eliminate an unknown, non-D1r target. In this report, we sought to better understand the proteomic consequences of microglial pruning in the NAc, and what the female pruning target might be. To do this, we inhibited microglial pruning in the NAc during each sex’s pruning period and collected tissue for mass spectrometry proteomic analysis and ELISA validation. We found that the proteomic consequences of inhibiting microglial pruning in the NAc were inversely proportional between the sexes, and a novel, female-specific pruning target may be Lynx1.

Please note, if this preprint will be pushed further to publication it will not be by me (AMK), as I am leaving academia. So, I’m going to write more conversationally.

## INTRODUCTION

Social behaviors are integral to health and wellness across the lifespan. Adolescence is a period of copious neural development, particularly in the ‘reward’ circuitry of the brain, and reward-related behavioral development, including social development [1-3]. One neurodevelopmental mechanism that appears to be common across brain regions and developmental periods is the requirement for synaptic pruning to produce mature neural communication and circuits [4-7]. In many cases, this is mediated via immune signaling: microglia, the resident immune cells of the brain, express complement receptor 3 (CR3/CD11b) which binds to its ligand C3, a phagocytic ‘tag’ that associates with weak or inactive synapses designated for pruning [8]. We published that microglia-C3-mediated synaptic pruning also occurs in the nucleus accumbens (NAc) reward region during adolescence to mediate social development in male and female rats [6]. However, both the adolescent stage in which microglial pruning occurred, and the synaptic pruning target, were sex specific. NAc pruning occurred between early and mid adolescence in male rats (postnatal day (P)30-40) to eliminate dopamine D1 receptors (D1rs), and between pre- and early adolescence in female rats (P20-30) to eliminate an unknown, non-D1r target [6]. Despite these differences, inhibiting microglial pruning in the NAc during each sex-specific pruning period with the highly specific CR3 antagonist, neutrophil inhibitory factor (NIF), acutely [6] and persistently [9] regulated social development in a similar manner in both sexes. Taken together these data suggest that a *convergent* developmental mechanism, NAc pruning, mediates sex-*divergent* neurochemical processing to induce sex-*convergent* social development. In this report, we sought to determine **(1)** whether the consequences of microglial pruning in the NAc, were, in fact, sexually divergent, and **(2)** what the female pruning target might be. To do this, we inhibited microglial pruning in the NAc during each sex’s pruning period and collected tissue for mass spectrometry proteomic analysis and ELISA validation. We found that the proteomic consequences of inhibiting microglial pruning in the NAc were inversely proportional between the sexes, and a novel, female-specific pruning target may be Lynx1.

## METHODS

### Animal Care

Adult male and female Sprague-Dawley rats were purchased to be breeding pairs (Harlan/Envigo). Litters were culled to a maximum of 12 pups between postnatal day (P)2-5, and pups were weaned into same-sex housing in pairs or triplets at P21. At least 3 litters were represented in each experimental group. For Novel Social Interaction and Social Choice tests, rats were purchased from Harlan/Envigo to act as conspecifics. Rats were housed in conventional cages on cellulose bedding with ad libitum access to food and water. Colonies were maintained on a 12:12 light:dark cycle (lights on a 07:00) in a temperature- (20-24°C) and humidity- (35-55 RH) controlled room. Cages were changed twice weekly. All experiments were approved by the Institutional Animal Care and Use Committee at Albany Medical College.

### Animal Model

Rats underwent surgical intervention at either P22 (females) or P30 (males) according to our previously published sex-specific NAc pruning periods [6]. Rats were anesthetized under isoflurane (2-3%) and fastened into the stereotactic apparatus. The scalp was cut midsagitally and holes drilled to target NAc bilaterally. NIF (5αg/αL) or Vehicle (sterile PBS) was infused into the NAc using a Hamilton syringe at 10° using sex-specific coordinate and volume parameters: AP +2.25mm, ML ±2.5mm, DV -5.75mm, 300nL for P30 males and AP +2.7mm, ML ±2.4 mm, DV -5.55mm, 250nL for P22 females. One depth was reached, the syringe remained in place for 1 min before infusion (50nL/min), and for 5 mins post-infusion. The wound was closed with surgical staples and treated topically with anti-bacterial ointment and Bupivicaine. Ketofen (5mg/kg) was administered subcutaneously after surgery, and once daily for two days after surgery.

### Sample Collection and Preparation

Rats were euthanized 8 days after surgery, a time point we have published that molecular and behavioral effects of NIF treatment are evident in both sexes [6], and perfused transcardially with saline. Brains were extracted, rapidly frozen, and 1mm punches were taken bilaterally from the NAc core. Tissue was homogenized in RIPA buffer with protease and phosphatase inhibitors. Homogenate from two sex- and manipulation-matched rats were pooled into one sample for proteomic analysis carried out by Creative Proteomics (https://www.creative-proteomics.com/).

### Mass Spectrometry (performed by Creative Proteomics)

200μg protein per sample was precipitated with cold acetone. Pellets were dissolved in 2M urea and denatured with 10mM DTT for 1hr at 56°C followed by alkylation with 50nM IAA for 1hr at room temperature in the dark. Ammonium bicarbonate was added to a final concentration of 50mM, pH 7.8, and trypsin was added to the sample for digestion for 15 hrs at 37°C. Peptides were purified with a C18 SPE column and lyophilized to near dryness. Peptides were resuspended in 20μL of 0.1% formic acid before LC-MS/MS analysis. 1μg of sample was loaded onto a nanoLC-MS/MS platform. The full scan was performed between 300-1650 m/z at a resolution of 60,000 at 200 m/z. The gain was set to 3e6. The MS/MS scan operated in Top 20 mode with 15,000 at 200 m/z resolution, 1e5 automatic gain control target, 19ms maximum injection time, 28% normalized collision energy, 1.4 Th isolation window, unassigned, 1, >6 charge state exclusion, and 30 sec dynamic exclusion. Raw MS files were aligned with the rat protein database using Maxquant (1.6.1.14). Protein modifications were carbamidomethylation (fixed), oxidation (variable), the enzyme specificity was set to trypsin, the maximum missed cleavage was set to 2, the precursor ion mass tolerance was set to 10 ppm, and the MS/MS tolerance was 0.5 Da.

### Data Analysis and Statistics

Data from Creative Proteomics were quality controlled in-house: any proteins that had more than one sample in which they were not detected (0 intensity) within the group of 3–4 total samples was excluded from analysis. In proteins with just one non-detected sampled, the average of the other 2-3 samples was used to replace the *t*-tests were performed between NIF and Vehicle groups for each sex. In the case of duplicate protein IDs, the ID corresponding to the smallest p-value was retained and the others were excluded from analysis. Protein groups that were significantly changed by NIF treatment (*p*<0.05, no correction for multiple comparisons) were assessed for global proteomic summary, including Pearson’s correlation of NIF-induced changes in all proteins between the sexes (*Fig. 1*). Benjamini-Hochberg analyses (FDR = 0.1) were performed to determine the proteins significantly changed by NIF treatment after correction for multiple comparisons (*Fig. 2*). Protein concentrations of remaining unpooled homogenate was determined with BCA assays following the manufacturer’s protocols. ELISAs were conducted per the manufacturer’s protocols by loading 2000μg total protein per sample. Lynx1 ELISA was purchased from mybiosource.com (#MBS9902329), NAChR ELISA was purchased from LSBio (#LS-F37436), and D1r ELISA was purchased from Novus Biologicals (NBP2-67935). ELISA data were analyzed with *t*-tests for each sex. There were no outliers detected via Grubb’s test (μ=0.05). For all analyses, statistical significance is defined as *p*<0.05. All statistics were performed in GraphPad Prism 9.5.1.

**Fig. 1:**
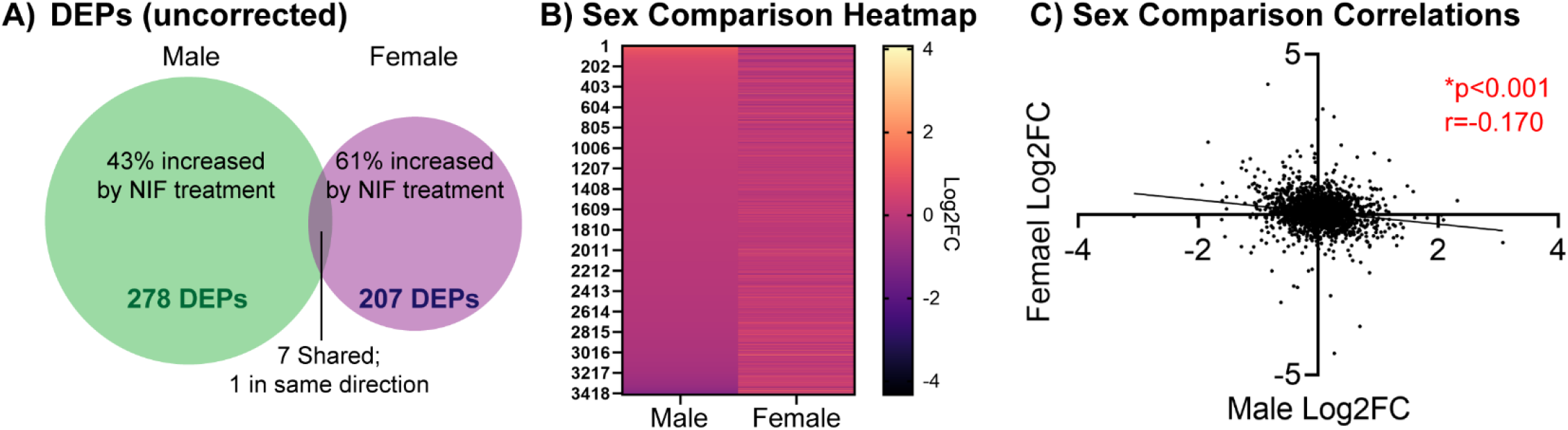
NIF-mediated inhibition of microglial synaptic pruning in the NAc induces inverse proteomic outcomes in male and female rats. Male and female rats were stereotactically injected with NIF or Vehicle bilaterally into the NAc at each sex’s respective pruning period (P30 in males, P22 in females). Eight days later, rats were euthanized, brains frozen, and NAc punches collected for proteomic analysis. Two sex- and treatment-matched rats’ NAc samples were pooled into one biological replicate to ensure enough protein yield. NAc samples were process via label-free nanoLC-MS/MS mass spectrometry. **(A)** After quality control, 3610 and 3687 total proteins were identified in male and female NAc, respectively, of which 278 and 207, respectively, were significantly changed (differentially expressed proteins; DEPs) at μ=0.05 uncorrected for multiple comparisons. Of these, 43% were upregulated by NIF-mediated inhibition of pruning in males, and 61% were upregulated in females. Only 7 total DEPs were shared between the sexes, and of the 7 only 1 DEP was regulated in the same direction (see **Supp. Table 1**). **(B)** Of all proteins identified, 3430 were shared between the sexes. Plotting the fold change of each protein in both sexes suggests that if a protein is upregulated by inhibiting NAc pruning in males, the same protein is likely to be downregulated by the same inhibition in females. **(C)** To assess this quantitatively, we examined the correlation in protein expression changes induced by NAc pruning inhibition between males and females, and found a significant negative correlation, indicating that our visual observation in *B* was the case. *n*=3-4 biological replicates/sex/condition. ^*^*p*<0.05.

**Fig. 2:**
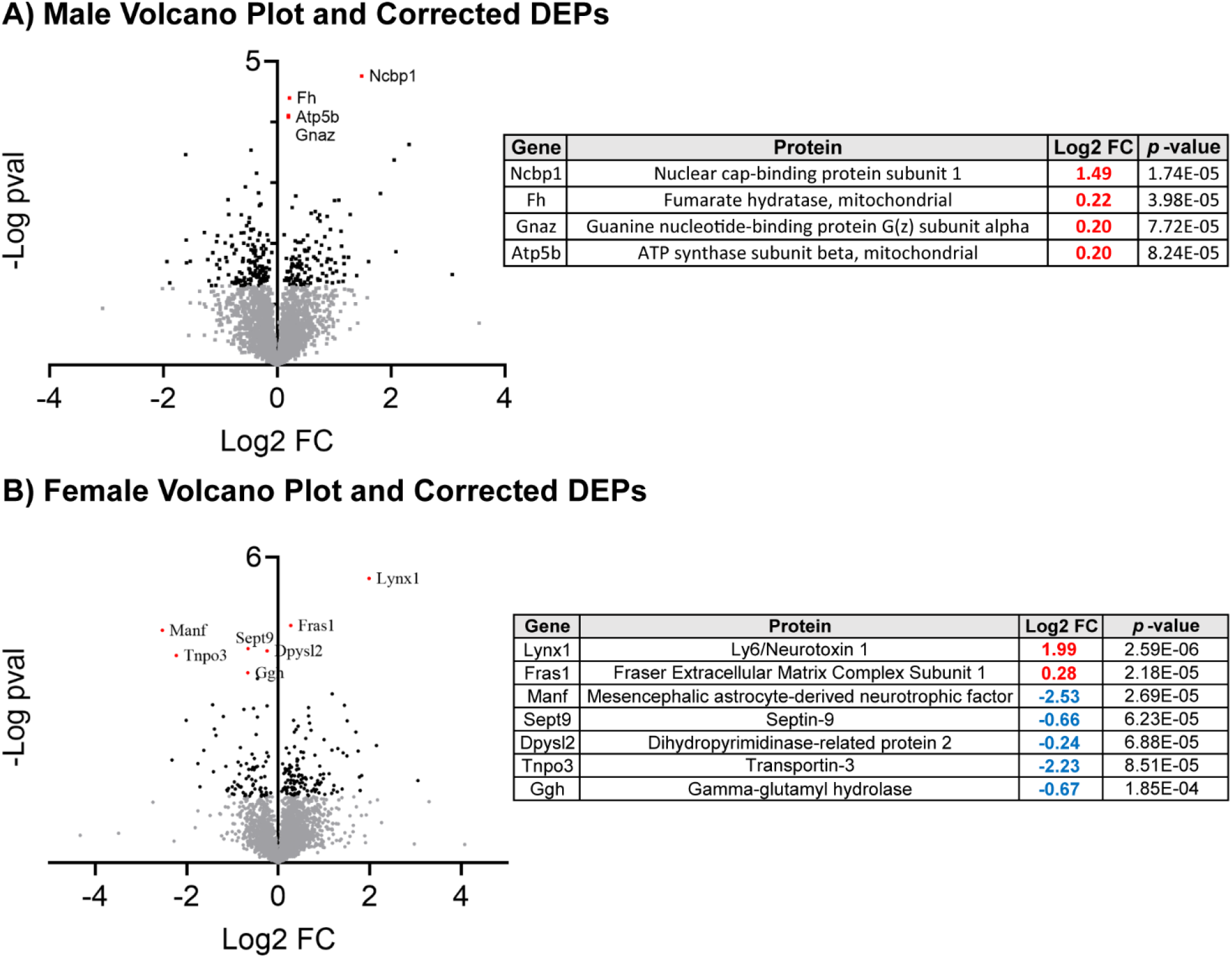
Sex-specific DEPs induced by NIF-mediated inhibition of microglial synaptic pruning in the NAc. Volcano plots depict each identified protein in each sex, with gray dots indicating non-significant (μ=0.05 uncorrected for multiple comparisons), black dots indicating significant DEPs (μ=0.05 uncorrected for multiple comparisons), and red, labeled dots indicating significant DEPs after correction for multiple comparisons (Benjamini-Hochberg; FDR=0.1). Proteins in Quadrant 1 are upregulated, and proteins in Quadrant II are downregulated. **(A)** In males, 4 DEPs passed multiple comparisons correction, which are identified in the table to the right. All four DEPs were upregulated (red FC value). **(B)** In females, 7 DEPs passed multiple comparisons correction, which are identified in the table to the right. Two DEPs were upregulated and the other 5 were downregulated (blue FC value). *n*=3-4 biological replicates/sex/condition.

## RESULTS

Because the male and female NAc pruning period is sex-specific [6], we performed surgeries to inhibit NAc pruning at different ages (P22 in females, P30 in males). Thus, we cannot directly compare the sexes.

However, we can compare patterns induced by inhibiting pruning in the NAc between the sexes. We will begin with a global summary of these patterns before examining unique outcomes in each sex.

### NIF-mediated inhibition of microglial synaptic pruning in the NAc induces inverse proteomic outcomes in male and female rats

In total, 3610 and 3687 total proteins were identified via label free mass spectrometry in NIF- and Vehicle-treated male and female NAc tissue, respectively. Of these, there were 278 differentially expressed proteins (DEPs; uncorrected for multiple comparisons) in males and 207 DEPs in females. Only 7 of these proteins were differentially expressed in both sexes, and of the 7 only one was regulated in the same direction (**Fig. 1A; Supp. Table 1**). A heatmap comparison of the NIF-induced fold change in all proteins suggested that male and female outcomes may be inversely related (**Fig. 1B**), which was confirmed with a significant negative correlation (*r*=-0.1697, *p*<0.001; **Fig. 1C**).

### NIF-mediated inhibition of microglial synaptic pruning in the NAc has sex-specific proteomic consequences

As discussed in Fig. 1, NIF-mediated inhibition of microglial pruning in the NAc regulated only 7 of over 200 DEPs in *both* sexes, 6 of which were regulated in different directions between the sexes. To better understand the unique consequences of NAc pruning in each sex, we conducted Benjamini-Hochberg adjustment for multiple comparisons, which revealed 4 DEPs in males and 7 DEPS in females that were still significantly regulated by NIF treatment. In males, these included the upregulation of Ncbp1, Fh, Gnaz, and Atp5b. In females, these included the upregulation of Lynx1 and Fras1 and the downregulation of Manf, Sept9, Dpysl2, Tnpo3, and Ggh. There was no meaningful pathway enrichment for either sex using either corrected or uncorrected DEP lists at the μ = 0.05 level (*data not shown*).

### Lynx1, but not NAChR, is regulated by microglia-mediated pruning in the NAc in females, but not males

When searching for a female-specific pruning target, we focused on proteins that were upregulated by NIF treatment (i.e., they would have been eliminated or downregulated by NAc pruning) and were synaptically-localized. In the female dataset, only Lynx1 fits this criteria via its close association with nicotinic acetylcholine receptors (NAChRs). Neither NAChr nor D1r, the known male pruning target [6], was identified in either sex via mass spectrometry. In fact, virtually no G-protein coupled receptors (GPCRs) were identified by mass spectrometry. We later learned that trypsin is not a good enzyme to for the identification of GPCRs using mass spectrometry, as there are few lysine residues in GPCRs for trypsin to cleave. Thus, in unpooled samples left from our mass spectrometry analyses, we quantified Lynx1, NAChR, and D1r via ELISA. Lynx1 was significantly regulated by NIF treatment in female, but not male NAc, but to our surprise the ELISA data showed a significant down-regulation (males: *t*(14)=1.225, p=0.241,; females: *t*(12)=2.647, p=0.021; **Fig. 3A**). There was no statistically significant regulation of NAChRs by NIF treatment in either sex (males: *t*(9)=2.138, p=0.061; females: *t*(10)=0.522, p=0.613; **Fig. 3B**). Finally, though we were running low on sample and thus did not have appropriate statistical power (*n*=3-5/sex/group), we did see an increase in D1r levels in NIF-treated males, but not females (**Fig. 3C**), consistent with our published results, indicating the NIF treatment was working as expected.

**Fig. 3:**
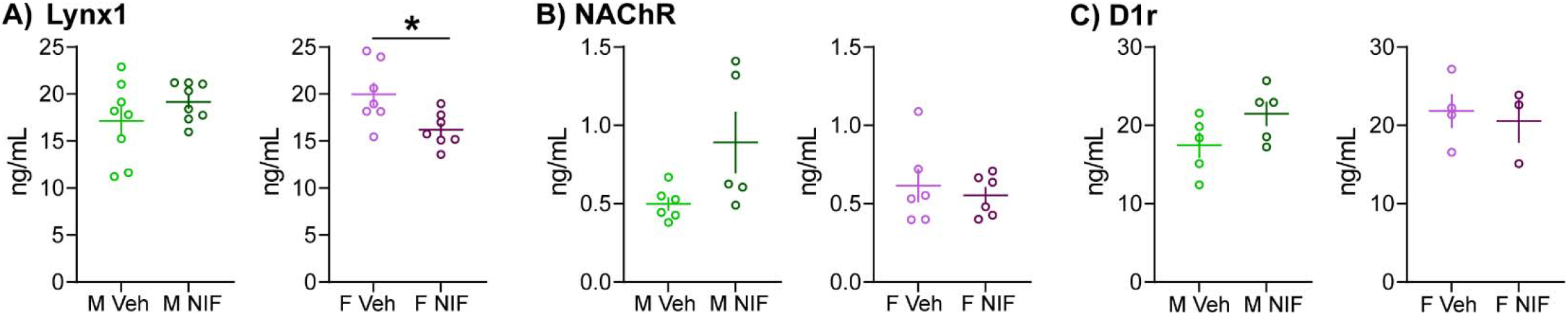
Lynx1, but not NAChR, is regulated by microglia-mediated pruning in the NAc in females, but not males. In remaining tissue (unpooled) that underwent proteomic analysis (*Figs. 1, 2*) we conducted ELISAs to validate potential female-specific pruning targets. **(A)** Inhibiting pruning in the NAc significantly decreased Lynx1 expression in females, but not males. *n*=7-8/sex/condition. **(B)** The Lynx1 target, NAChR, was not regulated by NIF treatment in either sex. *n*=7-8/sex/condition. **(C)** In remaining, but underpowered sample sizes (*n*=3-5/sex/condition), we observed the expected increase in D1r levels after inhibiting NAc pruning in males, but not females. This is consistent with our previously published results. In each histogram, horizontal lines are average and vertical lines are standard error of the mean. ^*^*p*<0.05.

## DISCUSSION

In this set of studies, we determined how microglia-mediated synaptic pruning in the NAc impacts the NAc proteomic landscape in male and female rats. We observed that despite manipulating the same developmental mechanism, synaptic pruning, the global proteomic outcome in the NAc was inversely related between males and females, and the sets of DEPs regulated by pruning were unique to each sex. Finally, we identify Lynx1 as a putative female-specific NAc pruning target.

### Same upstream developmental mechanism, different downstream molecular outcome

Sex differences of varying levels are now being reported in virtually every neural and behavioral outcome. Some sex differences are easily identifiable, e.g. a large change in behavior, while others are more subtle [10]. Our data indicate that *the same* developmental mechanism, C3-CR3 (microglial)-mediated synaptic pruning, is engaged in the adolescent NAc to mediate largely *the same* social development outcomes in both sexes. But the details are very different. Inverse, in fact! The same proteins that are upregulated by NAc pruning in males are more likely to be downregulated by NAc pruning in females, and vice versa. I don’t think that is a consequence of microglia just nibbling off different synaptic protein in males vs. females to induce downstream proteomic consequences in the pruning target’s respective signaling cascade. I think this is a mechanism that can sex-specifically reprogram the NAc, similar to the highly sex-divergent effects microglial pruning has on the amygdala [11] and preoptic area [12]. And if that is the case, it may be a mechanism that induces some of the sex differences that emerge during adolescence [3, 10]. Although NAc pruning does regulate social development, our data suggest that it does so in a more-or-less similar way in both sexes [6, 9]. We did observe that inhibiting pruning in the NAc during adolescence induced sex-specific expression of sociability towards a familiar (but not novel) social partner [9], but whether those subtle sex differences are meaningful remains unclear. Thus, it will be important to determine how NAc pruning during adolescence also impacts other reward-related behaviors to better understand the ways in which sex-specific NAc programing might be important.

### Lynx1 in brain and behavior

The proteomic data and ELISA validation identified Lynx1 as protein that was significantly regulated by microglial pruning in the NAc, specifically in females. However, (surprise!) the proteomic data indicated Lynx1 was increased by NIF treatment and the ELISA data indicated Lynx1 was decreased by NIF treatment. There are a number of reasons the data would not agree between these measurements. Mass spectrometry requires digestion of proteins into short peptides that are read and matched in a database. ELISAs measure native proteins using antibodies. It is very possible that a protein might be bound to other proteins in such a way that an antibody would not be able to bind. Lynx1 is an accessory protein that physically associates with NAChRs to modulate receptor function with very high efficiency [13]. If Lynx1 were upregulated in the NAc by inhibiting microglial pruning, then it stands to reason that more Lynx1 would be found bound to NAChRs in the tissue homogenate. More Lynx1 would equal more Lynx1 via mass spectrometry analysis, but more *bound* Lynx1 might equal less ‘free’, antibody-friendly Lynx1 for ELISAs. More validation will need to be performed, but it is worthwhile venturing down the Lynx1 as a female-specific pruning target path, as I think it could be quite impactful.

Lynx1 binding of NAChRs promotes receptor desensitization and slower recovery from desensitization [14]. Lynx1 has been demonstrated to alter NAChR subunit composition [15], but not necessarily total NAChR levels, which is supported by our data in which a change in Lynx1 levels was not accompanied by a change in NAChR levels (**Fig. 3**). Removing Lynx1 augments learning and memory [16], and interestingly also re-opens the critical period for visual plasticity [17], which is thought to be regulated by microglia-mediated synaptic pruning [4, 7]. Lynx1 removal is also associated with schizophrenia and autism spectrum disorders[18, 19], which are both linked to pruning abnormalities and social dysfunction (among many other things). Finally, Lynx1 removal can prolong adolescent plasticity required to establish social hierarchies after adolescent social isolation in mice [20]. Our data indicate that inhibiting microglial pruning in the female NAc increases Lynx1 expression, meaning that under normal developmental circumstances, microglia eliminate Lynx1 in the female NAc. Taken together with existing data, developmental elimination of Lynx1 should increase NAChR function in the NAc, augment NAc plasticity, and increase behavioral plasticity that is supported by NAc function – in females only. This will be important to thoroughly assess, as altered NAChR function in the NAc can impact multiple neurochemical systems [21] and may also point to a unique vulnerability (either mechanistically or in timing) of females at this adolescent stage to nicotine and ethanol [22] exposures.

## Acknowledgements

We thank Dr. Justin Bourgeois for assistance with tissue processing. This work was supported by the National Institutes of Health R01DA052889 and R03AG07011 to AMK and Albany Medical College Start-up funds to AMK.

## Author Declarations

AMK designed the experiments. JMK and IP performed the experiments. AMK analyzed the experiments. AMK wrote the manuscript. All authors edited the manuscript. The authors declare no conflicts of interest.

---

**Supp. Table 1:** DEPs regulated by NAc pruning shared between male and female rats. Only 1 of 7 DEPs regulated by NAc pruning, Ncbp1, was regulated in the same direction in male and female rats.

| Gene | Sex | Log2FC |
| --- | --- | --- |
| Slc4a1ap | F | **0.41172** |
|  | M | **-0.5752** |
| Cyb5r3 | F | **-0.4866** |
|  | M | **0.26419** |
| Rpl27 | F | **0.33966** |
|  | M | **-0.3207** |
| Ube2z | F | **0.77941** |
|  | M | **-0.2103** |
| Ncbp1 | F | **0.8934** |
|  | M | **1.4857** |
| Pccb | F | **0.35973** |
|  | M | **-0.3637** |

